# Modulatory effects of dynamic fMRI-based neurofeedback on emotion regulation networks in adolescent females

**DOI:** 10.1101/347971

**Authors:** Catharina Zich, Nicola Johnstone, Michael Lührs, Stephen Lisk, Simone P W Haller, Annalisa Lipp, Jennifer Y F Lau, Kathrin Cohen Kadosh

## Abstract

Research has shown that difficulties with emotion regulation abilities in childhood and adolescence increase the risk for developing symptoms of mental disorders, e.g anxiety. We investigated whether functional magnetic resonance imaging (fMRI)-based neurofeedback (NF) can modulate brain networks supporting emotion regulation abilities in adolescent females.

We performed three studies (total *N*=63). We first compared different NF implementations regarding their effectiveness of modulating prefrontal cortex (PFC)-amygdala functional connectivity (fc). Further we assessed the effects of fc-NF on neural measures, emotional/metacognitive measures and their associations. Finally, we probed the mechanism underlying fc-NF by examining concentrations of inhibitory and excitatory neurotransmitters.

Results showed that NF implementations differentially modulate PFC-amygdala fc. Using the most effective NF implementation we observed important relationships between neural and emotional/metacognitive measures, such as practice-related change in fc was related with change in thought control ability. Further, we found that the relationship between state anxiety prior to the MRI session and the effect of fc-NF was moderated by GABA concentrations in the PFC and anterior cingulate cortex.

To conclude, we were able to show that fc-NF can be used in adolescent females to shape neural and emotional/metacognitive measures underlying emotion regulation. We further show that neurotransmitter concentrations moderate fc-NF-effects.

## 1. INTRODUCTION

Adolescence is marked by a multitude of neural, emotional/metacognitive and behavioural changes, such as functional and structural maturation and improvements in cognitive abilities and socio-emotional behaviour (Blakemore, 2008; Burnett et al., 2011; Cohen Kadosh et al., 2013; Linscott and van Os, 2013). It has been suggested that these complex, transformational processes may shape poor emotion regulation abilities and increase the risk for developing mental disorders (Haller et al., 2015, 2013; Keshavan et al., 2014; Paus et al., 2008). Anxiety is one of the most common and impairing mental disorders in adolescence. Youth with anxiety experience impairing levels of fears and worries that they are unable to regulate, which can impact friendships, school performance, and mark the beginning of long-term mental health difficulties (Beddington et al., 2008; Pine et al., 2001; Trentacosta and Fine, 2010). Gaining a better understanding of the mechanisms underlying the successful regulation of emotions during this unique developmental period therefore represents an important step towards devising efficient, and targeted strategies for early support and intervention.

In the human brain, the regulation of emotions relies on a network of brain regions, comprising the prefrontal cortex (PFC) and the amygdala, as well as additional regions in the orbitofrontal, frontal and cingulate cortex (Kohn et al., 2014; Ochsner and Gross, 2005). A particular focus has been placed on the PFC-amygdala relationship, a relationship that manifests in both structural and functional connections (Banks et al., 2007; Davidson et al., 2000; Kim et al., 2003; Kim and Whalen, 2009; Kohn et al., 2014; Quirk and Beer, 2006). With regard to the regulatory aspect of the PFC-amygdala connection, the functional connectivity (fc) between the two regions has been theorized to reflect top-down PFC regulation of amygdala reactivity (Hare et al., 2008; Hariri et al., 2003; Kim et al., 2003; Pezawas et al., 2005), with increased PFC responses leading to decreases in amygdala activation, i.e. negative fc (Kim et al., 2011). Interestingly, this fc pattern only emerges during adolescence (Dougherty et al., 2015; Gee et al., 2013; Wu et al., 2016), with research showing that children at the beginning of adolescence (up until around 10 years of age) exhibit predominantly positive fc, rather than negative fc ((Dumontheil et al., 2012, 2010; Silvers et al., 2012), also reviewed in (Ahmed et al., 2015)). What is less clear however is whether we can intervene to actively shape fc patterns in the developing brain to improve emotional/metacognitive abilities and thereby affect behaviour.

The current study aimed to address this question by using real-time functional magnetic resonance imaging (fMRI)-based neurofeedback (NF). NF provides the user with real-time information of one’s own brain activity, and it has already been shown to be a promising intervention tool to (re-)shape neural activity in a number of psychiatric and neurological diseases (Linden, 2012; Lubianiker et al., 2019). In the case of emotion regulation, NF can aid the modulation of networks underlying this process, such as PFC-amygdala fc. Previous studies suggest that fMRI-based NF represents a promising approach to train individuals in the self-modulation of brain regions or networks (Johnston et al., 2010; Koush et al., 2017, 2013; Scharnowski and Weiskopf, 2015; Weiskopf et al., 2004; Zilverstand et al., 2014), also in the emotion processing domain (adults: (Johnston et al., 2010; Koush et al., 2017; Paret et al., 2014; Zotev et al., 2013); children: (Cohen Kadosh et al., 2016)).

Building upon these studies, we conducted a set of three experiments that investigated whether fMRI-based fc-NF of PFC-amygdala fc can be used to modulate neural measures (e.g. change in fc) and emotional/metacognitive measures (i.e. self-reported measures of anxiety and thought control ability) relevant for emotion regulation in adolescent females. To achieve this overall objective, we first contrasted three different NF implementations with regard to their effectiveness in modulating PFC-amygdala fc (**Experiment 1**). The most effective NF implementation was then used to assess the effects of one session fc-NF on neural measures and emotional/metacognitive measures in a larger sample (**Experiment 2**). Finally, we assessed the effects of a longer fc-NF block in order to further enhance the effectiveness of the fc-NF and explored how neurotransmitter concentrations in relevant regions influence the observed effects (**Experiment 3**). Regarding the latter, we used proton magnetic resonance spectroscopy (^1^H-MRS) to extract the neurotransmitter profile from two voxels of interest (VOI), the PFC and the anterior cingulate cortex (ACC). We choose the PFC because of the key role of γ-aminobutyric acid (GABA)-ergic neurotransmission within the PFC in balancing amygdala activity (Constantinidis et al., 2002). Further, we choose the ACC because of its centrality in the reward processing network, especially for explicit processing of reward, such as NF (Emmert et al., 2016; Sitaram et al., 2016). For both of these VOIs we quantified GABA and glutamate, the major inhibitory and excitatory neurotransmitters in the human brain.

## 2. MATERIALS AND METHODS

### 2.1. Experiment 1

#### Participants

18 female adolescent participants (*M* = 14.83 years; *SD* = 0.99 years) were recruited from local schools in the Oxfordshire/Gloucestershire area. The single-sex approach allowed us to minimize variation introduced by differences in hormonal development and puberty (Goddings et al., 2014, 2012). All participants had normal or corrected-to-normal vision and reported no history of neurological and psychiatric disorders (determined via self-report). Informed written consent was obtained from the primary caregiver and informed written assent was obtained from the adolescent. Participants received an Amazon voucher (£20) for their participation. The study was approved by the Central Oxfordshire Ethics Committee (MSD-IDREC-C2-2015-023) and conducted in accordance with the Declaration of Helsinki. This work was registered as preclinical trial (ClinicalTrials.gov Identifier: NCT02463136).

#### Self-report questionnaires

Immediately prior to the MRI session, participants completed several self-report questionnaires, which will be referred to as *emotional/metacognitive measures*. Specifically, we assessed psychological variables using the Emotion Regulation Questionnaire (CERQ) (Derryberry and Reed, 2002), the Mood and Feelings Questionnaire (MFQ) (Angold et al., 1995), Social Anxiety Scale for Adolescents (SAS-S) (La Greca and Lopez, 1998), the State-Trait Anxiety Inventory (STAI-T, STAI-S) (Spielberger et al., 1999), the Thought Control Questionnaire (TCQ) (Wells and Davies, 1994) and the Thought Control Ability Questionnaire (TCAQ) (Luciano et al., 2005). These are established self-report measures, which are frequently used in psychological testing. A subset of the questionnaires (i.e. CERQ, MFQ, SATI-S, TCQ, TCAQ) were repeated after the MRI session in order to assess changes. Participants also completed a Demographic and Health Questionnaire and the Wechsler Abbreviated Scale of Intelligence (Wechsler, 2011).

#### MRI data acquisition

MRI data acquisition was performed on a 3 T Siemens MAGNETOM Prisma MRI scanner (Siemens AG, Erlangen, Germany) using a standard 32-channel head matrix coil. First, a high-resolution structural scan was acquired, which was followed by functional imaging during the localizer task and the NF task (see **Supplementary Methods** for details on the MRI sequences).

#### Localizer task

A modified version of the social scenes task (Haller et al., 2016) was used to identify the NF regions of interest (ROI), as it was expected that this task activates the key regions involved in cognitive and emotional appraisal and reappraisal. The localizer task lasted 8.9 min. and comprised 30 trials. Each trial started with a social scene presented for four volumes (3.73 s). Scenes depicted negative, rejecting social situations viewed from the perspective of a female protagonist depicted from the back. Participants were instructed to interpret the scene freely (appraisal). This was followed by a positively valanced interpretative statement (4 volumes, 3.73 s), after which the same scene was shown again for four volumes (3.73 s). For the duration of the second presentation, participants were instructed to reappraise the scene based on an interpretative statement (reappraisal). Finally, participants were asked to rate how much they were able to change, i.e. reappraise, their thoughts and feelings from the first to the second presentation of the scene. Participants indicated their perceived change on a Likert scale ranging from no change (1) to much change (4) via button press on each trial. A fixation cross was presented for one volume between two trials. Stimulus presentation was controlled via BrainStim 1.1.0.1 (open source stimulation software, Maastricht University, http://svengijsen.github.io/BrainStim/). Participants were encouraged to use these emotional reappraisal strategies during the NF tasks.

#### Neurofeedback task

The NF task consisted of four identical runs lasting 4.8 min each. Stimulus presentation was controlled with BrainStim 1.1.0.1. Each run started with a fixation cross that was displayed for 20 volumes (18.66 s), followed by seven fc-NF mini-blocks (20 volumes each) and seven no-NF mini-blocks (20 volumes each) that were presented in alternating order, with the start condition being randomized and counterbalanced across individuals. During both fc-NF and no-NF mini-blocks a ten-segment thermometer was presented at the centre of the screen on a dark grey background. During the no-NF mini-blocks, the ‘temperature’ of the thermometer was frozen at the sixth segment (i.e. 6/10) throughout the mini-block (**Fig. 1a**). During the fc-NF mini-blocks, the thermometer featured a green frame and the ‘temperature’ was a direct reflection of PFC-amygdala fc and was updates with every TR (see section ‘Online MRI data analysis’ for more details). All participants were asked to up-regulate the thermometer by controlling their thoughts and feelings and by revisiting emotional reappraisal strategies as practised in the localizer task (see **Supplementary Methods** for more details on the instructions).

**Fig. 1.**
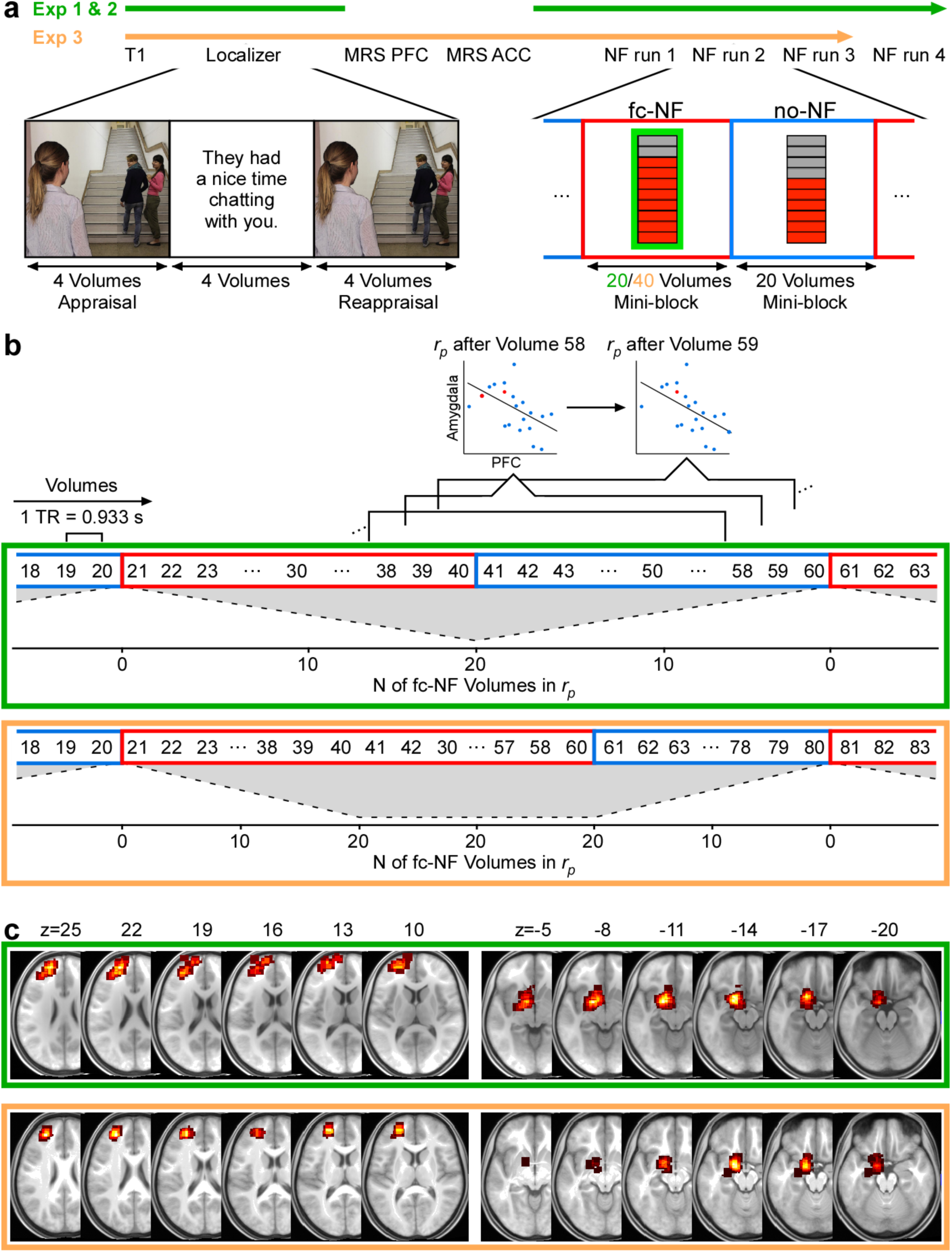
Schematic representation of the experimental procedures (**Experiment 1** and **2** in green and **Experiment 3** in orange). **(a)** Experimental timelines. To activate the key regions involved in emotion regulation participants performed the social scenes task (localizer). Based on an individual’s localizer activity, we then defined three regions of interests (ROIs): left prefrontal cortex (PFC), left amygdala, left corticospinal tract (CST). Each NF run comprised fc-NF mini-blocks (20 volumes in **Experiments 1 and 2**; 40 volumes in **Experiment 3**) and no-NF mini-blocks (20 volumes), presented in alternating order. **(b)** Detailed view of the procedure during the NF task. NF was based on partial correlations *r*_*p*_ between the PFC and amygdala activity, while controlling for CST ‘activity’, whereby partial correlations were obtained from a moving window comprising 20 subsequent volumes, which was updated with every incoming volume. Thus, for **Experiments 1** and **2**, the ratio of fc-NF : no-NF volumes in the correlation window is 1:19 at the first volume of the mini-block, 2:18 at the second volume of the mini-block and 20:0 at the last volume of the mini-block, and vice versa for no-NF mini-blocks. Note that due to the extended window length for the fc-NF mini-blocks in **Experiment 3**, the ratio of fc-NF : no-NF volumes in the correlation window is 20:0 for twenty volumes. Number of fc-NF volumes in the correlation window (N of fc-NF Volumes in *r*_*p*_) is illustrated in form of a grey shape. **(c)** Probabilistic maps of subject-specific ROIs (i.e. PFC and amygdala) superimposed on the groups’ average structural image in MNI space. The respective centre of the PFC probabilistic maps (Experiment 1 and 2: x=-20, y=50, z=24; Experiment 3: x=-26, y=47, z=22) are classified as Brodmann area 10 within the left middle frontal gyrus. The respective centre of the amygdala probabilistic maps (Experiment 1 and 2: x=-17, y=-1, z=-14; Experiment 3: x=-16, y=-3, z=-15) are classified as left amygdala (Eickhoff et al., 2005).

We evaluated three different NF implementations with regard to their effectiveness of modulating PFC-amygdala fc (*N* = 6 per implementation). The three different NF implementations only differed in the upper value of the thermometer (the lower value of the thermometer was set to zero). The upper value of the thermometer was set to −1 (maximal negative correlation) for the *negative NF implementation*, which reinforced negative fc; to −0.3^†^ for the *weighted negative NF implementation*, which reinforced negative fc; and to +1 (maximal positive correlation) for the *positive NF implementation*, which reinforced positive fc. To illustrate this: if the thermometer changes from 2/10 to 3/10 segments, this corresponds to a more negative correlation by 0.1 for the negative NF implementation, a more negative correlation by 0.03 for the weighted negative NF implementation, and a change towards a more positive correlation by 0.1 for the positive NF implementation.

#### Online MRI data analysis

To enable real-time fMRI-based NF, MR images were passed from the MRI console computer to the real-time computer via a direct TCP/IP network link using the Server Message Block (SMB) network layer. Following completion of the structural scan, anatomical images were processed using BrainVoyager QX 2.8.2 (Brain Innovation, Maastricht, The Netherlands). Each 3D volume was corrected for B0 inhomogeneities (4-cycle bias field estimation), followed by brain extraction from the skull and separating bone and cerebro-spinal fluid (2-cycle iteration).

Functional images obtained during the localizer and the NF runs were processed in real-time using Turbo-BrainVoyager 3.2 (Brain Innovation, Maastricht, The Netherlands). To correct head motion, each volume was realigned to the first volume of the localizer. Each realigned volume was smoothed with a three-dimensional Gaussian kernel of 8 mm full-width-half-maximum. After completion of the localizer, three ROIs (voxel size = 12 mm^3^, 6 × 6 × 6) were selected. To this end, GLM t-statistics of the brain activity during the localizer task, i.e. the sum of the three contrasts: appraisal > fixation, reappraisal > fixation, reappraisal > appraisal (threshold *t* = 3), was projected onto the processed structural scan. The local maximum of the t-statistics within the left dorsolateral and medial PFC constituted the centre of the PFC ROI. Similarly, the local maximum of the t-statistics within the left amygdala constituted the centre of the amygdala ROI. A ROI in the left corticospinal tract (CST) served as control ROI^‡^. Probability maps of the selected ROIs across subjects are depicted in **Fig. 1c.**

During the NF task, PFC-amygdala fc was calculated in real-time. PFC-amygdala fc was defined as the partial correlation *r*_*p*_ between PFC and amygdala activity, while controlling for CST ‘activity’. As shown previously, partial correlation analysis can be used to quantify fc between areas while controlling for noise from a task-unrelated region (Dawson et al., 2016). Partial correlations were based on a moving window, which was updated with every incoming volume. The length of the correlation window was 20 volumes (see **Supplementary Discussion**). Therefore, the ratio of fc-NF to no-NF volumes in the correlation window changed with every incoming TR (**Fig. 1b**). To illustrate this: for the fc-NF mini-blocks, the ratio of fc-NF : no-NF volumes in the correlation window was 1:19 at the first volume of the mini-block, 2:18 at the second volume of the mini-block and 20:0 at the last volume of the mini-block, and vice versa for no-NF mini-blocks. Calculations were performed using a custom-made plugin for Turbo-BrainVoyager, which also provided a direct TCP/IP based link between the real-time analysis software and the stimulus application BrainStim. PFC-amygdala fc was displayed via thermometer during the fc-NF mini-blocks.

#### Offline MRI data analysis

We also we analysed the MRI data offline. While online and offline analysis of the MRI data follow a similar processing pipeline, offline algorithms tend to be more robust. Offline analysis of the MRI data was performed using SPM12 (FIL, Wellcome Trust Centre for Neuroimage, UCL, London, UK). Here head motion was corrected by realigning the functional time series of the localizer or the NF runs to its first volume. Due to motion artefacts (i.e. motion exceeded 3 mm on any axis) three individuals (i.e. one individual per group) were excluded from all analyses. Each individual’s structural image was registered to their mean functional image and segmented, in order to normalize structural and functional images to the Montreal Neurological Institute (MNI) template. Finally, normalized functional images were smoothed with a three-dimensional Gaussian kernel of 8 mm full-width-half-maximum. To increase the consistency between online and offline analysis we used the same ROIs. Therefore, the ROIs defined in Turbo-BrainVoyager during the MRI session were transformed from DICOM to NIFTI format, from radiological to neurological convention and from voxel to mm space. Subsequently, ROIs were normalized to the MNI template by applying the same transformation matrix used for the subject-specific normalization of structural and functional images. Time series were extracted from the three ROIs using MarsBaR 0.44 (Brett et al., 2002) and PFC-amygdala fc calculated using the same procedure as during real-time NF, i.e. partial correlation *r*_*p*_ between PFC and amygdala activity, while controlling for CST ‘activity’.

#### Statistical analysis

To identify the most effective NF implementation we evaluated the difference between no-NF and fc-NF for each NF implementation. The primary measure for this comparison constitutes PFC-amygdala fc, i.e. partial correlations *r*_*p*_. Fc was averaged at each of the 20 volumes (i.e. length of a mini-block) across the seven mini-blocks within one condition (fc-NF, no-NF), NF runs (1, 2, 3, 4) and individuals within one NF implementation (negative NF implementation, weighted negative NF implementation, positive NF implementation). The relationship of fc-NF and fc was assessed by correlating (Pearson’s *r* correlations as implemented in SPSS version 25; SPSS Inc, Chicago, IL, USA) the average fc at each of the 20 volumes with the number of fc-NF volumes within the correlation window (*N* = 40, i.e. 20 volumes per mini-block). To statistically compare these Pearson’s *r* correlations between the different NF implementations, Fisher’s z-transformation was applied to each correlation coefficient, resulting in normally distributed values *r’* with standard errors *s*_*r*_*’*. The null hypotheses (*r’*1-*r’*2 = 0) were tested in R(psych) (Revelle, 2015) using Student *t* test (Howell, 2011).

The same analysis was conducted to assess the frequency of negative partial correlations (secondary measure) and the frequency of scenarios describing different patterns of activation changes (**Supplementary Fig. 5**).

### 2.2. Experiment 2

This study comprises data from 30 adolescent females (*M* = 15.20 years; *SD* = 1.10 years; 25 naïve individuals and the 5 individuals tested using the weighted negative NF implementation in **Experiment 1**). The recruitment procedure was identical to **Experiment 1**. The ‘Self-report questionnaires’, ‘MRI data acquisition’, ‘Localizer task’, ‘Neurofeedback task’ as well as the ‘Online MRI data analysis’ and ‘Offline MRI data analysis’ were identical to **Experiment 1**. However, all individuals received NF using the weighted negative NF implementation. Due to motion artefacts (motion exceeded 3 mm on any axis) datasets from three individuals were excluded from all analyses.

#### Statistical analysis

Firstly, we performed the same analysis as in **Experiment 1**.

We extended this by investigating practice-related change in emotional/metacognitive measures and neural measures. *Practice-related change in emotional/metacognitive measures* was defined as difference between emotional/metacognitive measures obtained before and after the MRI session and assessed using paired samples *t* test (SPSS). Practice-related change in neural measures, i.e. *practice-related change in fc*, was defined as the slope of the linear regression of the total fc across runs (**Supplementary Fig. 4a**) and was tested using a one-sample *t* test (SPSS).

Moreover, we performed correlation analyses between different emotional/metacognitive and neural measures. Relevant emotional/metacognitive measures were practice-related change in emotional/metacognitive measures (as defined in the previous paragraph) and *initial emotional/metacognitive measures*, i.e. measures obtained before the MRI session. Relevant neural measures were practice-related change in fc (as defined in the previous paragraph), initial fc and fc-NF-effect. *Initial fc* was defined as the average fc of the first two mini-blocks (**Supplementary Fig. 4b**). *Fc-NF-effect* was defined as the difference between no-NF and fc-NF (**Supplementary Fig. 4c**). Correlation analysis was performed using Bayesian inference (JASP, JASP Team 2019, version 0.8.1.2) with default priors. Results are reported for the whole sample and after outlier removal. Outliers were identified for each correlation separately by bootstrapping the Mahalanobis distance (Schwarzkopf et al., 2012).

Missing values analysis revealed that 2.56% of the self-report questionnaire data were missing. We performed Little’s test of Missing Completely at Random (MCAR) (Little, 1988), as implemented in SPSS. MCAR was not significant (*X*^*2*^_(35, *N* = 27)_ = 41.74, *p* = .201), i.e. there is no evidence to suggest that the data were not MCAR. As such, pairwise deletion was used in the statistical analysis.

### 2.3. Experiment 3

20 naïve female adolescent participants (*M* age = 15.85 years; *SD* = 1.04 years) were tested using the weighted negative NF implementation as described in **Experiment 1**. The recruitment procedure was identical to **Experiments 1** and **2**.

The ‘Self-report questionnaires’ were identical to **Experiments 1** and **2**. ‘MRI data acquisition’ was similar to **Experiments 1** and **2**, the only modification beeing the number of volumes during the localizer and the NF runs (see **Supplementary Methods**). The ‘Localizer task’ was similar to **Experiments 1** and **2**, however, only half of the trials were performed (i.e. 15 trials, 4.7 min.).

The ‘Neurofeedback task’ was similar to **Experiments 1 and 2**, with one critical variation: we lengthened the fc-NF mini-blocks to 40 volumes while the no-NF mini-blocks remained 20 volumes. We collected data from 3 identical runs, each comprising five fc-NF mini-blocks and five no-NF mini-blocks. The ‘Online MRI data analysis’ and ‘Offline MRI data analysis’ was similar to **Experiments 2** taking into account the change from 20 to 40 volumes per fc-NF mini-block. Moreover, we collected MRS data from two VOIs, the PFC and the ACC. See **Supplementary Methods** for details on MRS data acquisition. Due to motion artefacts (motion exceeded 3 mm on any axis) the dataset from one individual was excluded from all analyses.

#### MRS analysis

MRS post-processing was performed using the MATLAB (Mathworks, Natick, MA) toolbox MRspa (version 1.5f, https://www.cmrr.umn.edu/downloads/mrspa/). Motion corrupted spectral averages were removed and frequency and phase drifts corrected before spectral averaging. Averaged spectra were quantified in LCmodel (Provencher, 2001, 1993) in reference to a simulated basis set. GABA and glutamate were quantified in reference to total Creatine (Creatine + Phosphocreatine, t/CrPCr) for each VOI. Cramér–Rao lower bounds were <20% for GABA (t/CrPr) indicating reliable estimates extracted from spectra for all but two values in the PFC which were 22% and 27%. Both were retained for analysis. GABA (t/CrPr) concentrations were similar in each VOI (*t*_(18)_ = 0.89, *p* = .386; ACC: *M* = 5.811%, *SD* = 1.40%, PFC: *M* = 5.29%, *SD* = 2.25%). Cramér–Rao lower bounds were all <20% for Glutamate (t/CrPr) indicating reliable estimates extracted from spectra. Glutamate (t/CrPr) concentrations differed by VOI (*t*_(18)_ = 4.85, *p* < .001), with estimates higher in the ACC (*M* = 10.57%, *SD* = 0.76%), than the PFC (*M* = 9.34%, *SD* = 0.75%).

#### Statistical analysis

We performed the same statistical analysis as in **Experiment 1** and **2**. In addition, we conducted a moderation analysis following mean centering using the SPSS add-on tool PROCESS macro (version 3.3) (Hayes, 2018). Significant interactions were followed-up with simple slope analysis.

## 3. RESULTS

### 3.1. Experiment 1

The aim of **Experiment 1** was to compare different NF implementations regarding their effectiveness of modulating PFC-amygdala fc. The primary measure for this comparison, i.e. PFC-amygdala fc, is illustrated in **Supplementary Fig. 1a**.

The relationship of fc-NF and fc was assessed by correlating the average fc at each of the 20 volumes with the number of fc-NF volumes within the respective correlation window (**Supplementary Fig. 1b**). For the weighted negative NF implementation, a significant negative relationship was found (*r*_(38)_ = −0.72, *p* < 0.001), i.e. the more fc-NF volumes were included in the correlation window, the less positive the resulting partial correlation. The opposite relationship was found for the negative NF implementation (*r*_(38)_ = 0.37, *p* = 0.02), i.e. the more fc-NF volumes were included in the correlation window, the more positive the resulting partial correlation. No significant relationship was observed for the positive NF implementation (*r*_(38)_ = 0.22, *p* = 0.16).

Finally, the relationship between the average fc at each volume and the number of fc-NF volumes within the correlation window observed for the weighted negative fc-NF implementation was significantly different from the other two implementations (both *p*’s < 0.001), which did not differ from each other (*p* = 0.48). The picture is similar for the secondary measure, i.e. the frequency of negative partial correlations (see **Supplementary Results, Supplementary Fig. 2, Supplementary Fig. 3** for details).

In sum, these findings demonstrate the superiority of the weighted negative NF implementation in modulating PFC-amygdala fc towards more negative fc^§^.

### 3.2. Experiment 2

The aim of **Experiment 2** was to assess the effects of one session fc-NF (weighted negative NF implementation) on neural measures, emotional/metacognitive measures and their associations in a larger sample. The fc-NF-effects observed in **Experiment 1** were replicated, that is, the number of fc-NF volumes within the correlation window was negatively related with the average fc at each volume (**Fig. 2a**) and positively related with the frequency of negative partial correlations at each volume (**Fig. 2b**).

**Fig. 2.**
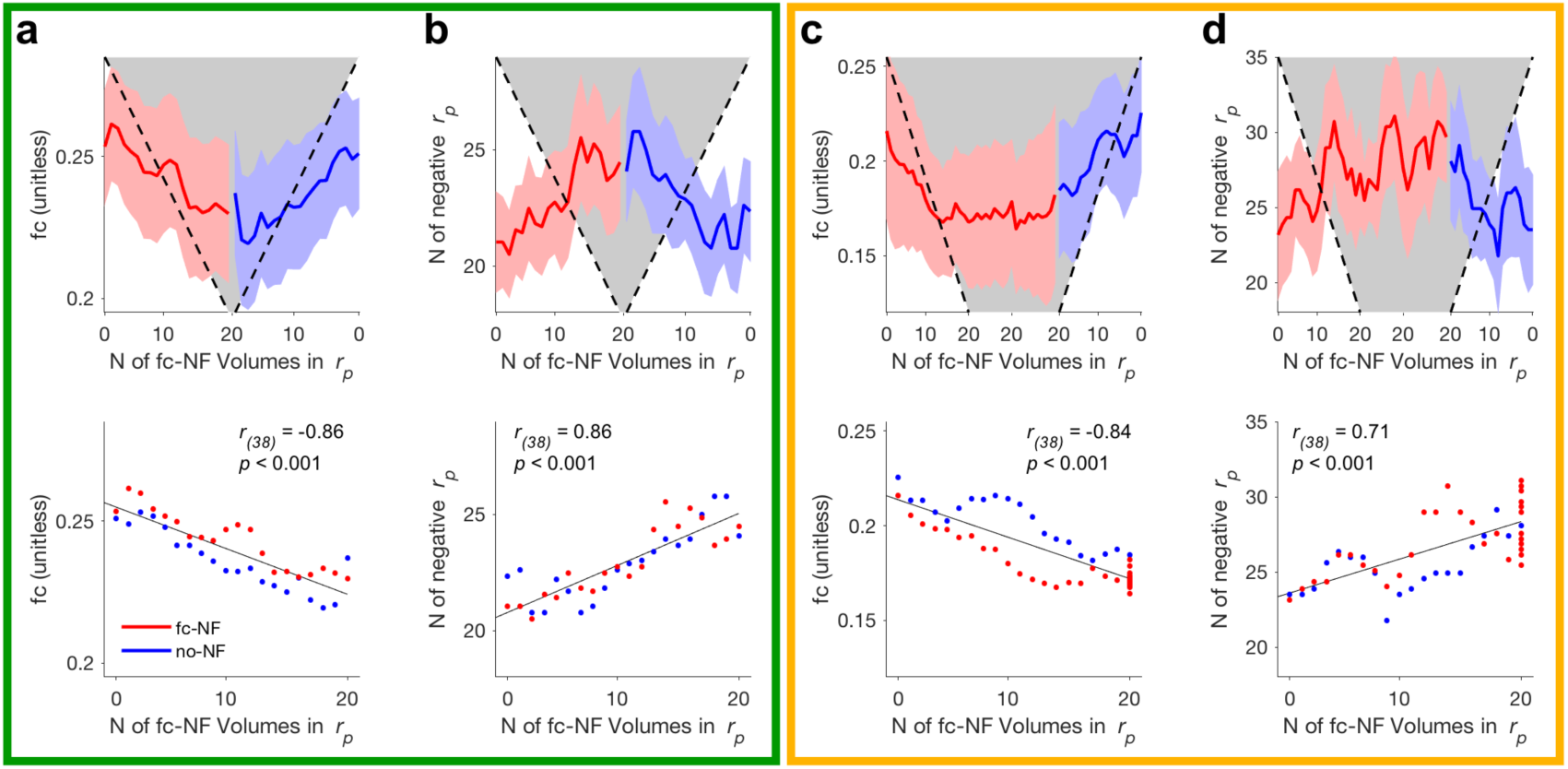
Difference in average functional connectivity (fc) and frequency of negative partial correlations (N of negative *r*_*p*_) between fc-NF (red) and no-NF (blue) for **Experiment 2** (green frame) and **Experiment 3** (orange frame). **(a, c)** Top: Means and standard error for average fc, i.e. partial correlations *r*_*p*_, for fc-NF and no-NF at each volume. Fc is averaged across mini-blocks within one condition (fc-NF, no-NF), NF runs and individuals. Number of fc-NF volumes in the correlation window (N of fc-NF Volumes in *r*_*p*_) is illustrated in form of a grey shape. Bottom: Relationship between average fc at each volume and the number of fc-NF volumes within the correlation window. **(b, d)** Same as (a, c), but for the frequency of negative partial correlations.

We further investigated practice-related change on emotional/metacognitive measures and neural measures for the whole sample. At emotional/metacognitive level no significant changes between measures obtained before and after the MRI session were observed (**Supplementary Table 2**). Similarly, at neural level no significant practice-related change in fc could be observed (*M* = .005, *SD* = .035, *p* > .01). Thirteen individuals showed a negative slope (i.e. change in the desired direction, *M* = -.023, *SD* = .019), fourteen individuals exhibited a positive slope (*M* = .031, *SD* = .025). Note that the emotional/metacognitive measures were comparable between these two subsamples (see **Supplementary Table 3** and **Supplementary Table 4**).

#### Correlation analysis

We first investigated whether practice-related change in emotional/metacognitive measures and practice-related change in fc were related. We found that practice-related change in fc was negatively related with change in TCAQ (**Fig. 3a** top; *r*_(21)_ = -.37, B_10_ = 1.03; after the removal of one outlier: *r*_(20)_ = -.58, B_10_ = 11.00). Hence, the larger the practice-related change towards negative fc, the better the thought control ability after the MRI session, when compared to the before the MRI session. Moreover, we were interested whether, neural measures could be predicted by initial emotional/metacognitive measures, i.e. measures obtained before the MRI session. Correlation analysis revealed that initial fc was positively related with STAI-T (**Fig. 3b** top; *r*_(25)_ = .36, B_10_ = 1.17; after the removal of two outliers: *r*_(23)_ = .59, B_10_ = 23.66) and negatively related with thought control ability (TCAQ: **Fig. 3c** top; *r*_(21)_ = -.36, B_10_ = 1.03; after the removal of two outliers: *r* _(19)_ = -.66, B_10_ = 35.01). In other words, initial fc was more negative in individuals with lower trait anxiety and better ability to control thoughts before the MRI session. Further, we found that STAI-S and fc-NF-effect were negatively related (**Fig. 3d** top; *r*_(22)_ = -.41, B_10_ = 1.67; after the removal of one outlier: *r*_(21)_ = -.55, B_10_ = 7.78). This suggests that individuals with lower state anxiety before the MRI session showed a larger difference between fc-NF and no-NF than individuals with higher state anxiety.

**Fig. 3.**
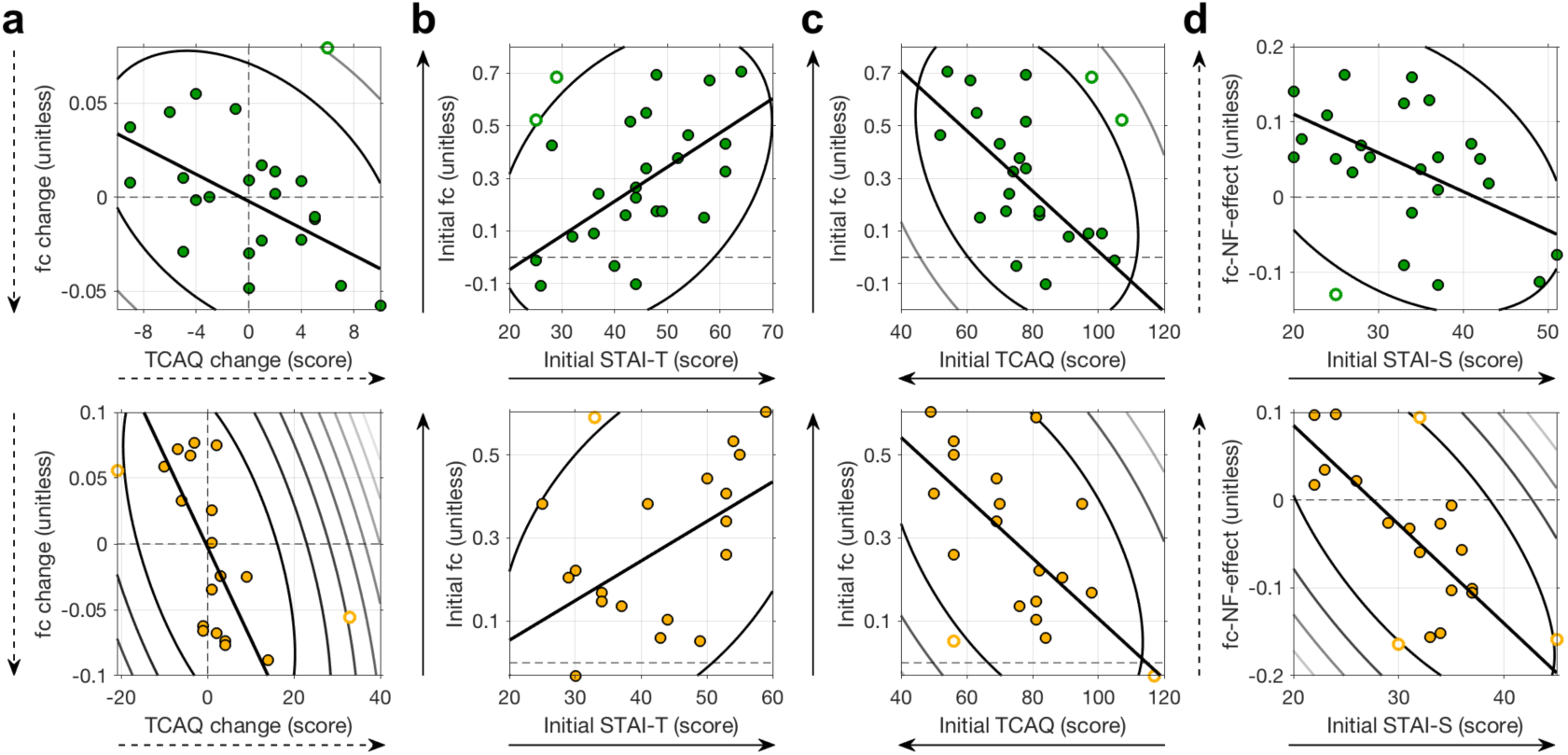
Associations between the emotional/metacognitive and neural measures for **Experiment 2** (top, green) and **Experiment 3** (bottom, orange). (**a**) Association between change in fc and change in thought control ability (**b**,**c**) Association between the initial fc and trait anxiety (STAI-T) as well as thought control ability (TCAQ) obtained before the MRI session. (**d**) Association between fc-NF-effect and state anxiety (STAI-S) obtained before the MRI session. Counter lines represent the bootstrapped Mahalanobis distance from the bivariate mean in steps of six squared units. Open circles represent outliers and the solid line is the regression over the data after outlier removal. Direction of solid arrows indicate levels related with higher anxiety. Direction of dashed arrows indicate the desired direction of practice-related change and fc-NF-effect.

### 3.3. Experiment 3

The aim of **Experiment 3** was three-fold: to assess the effect of longer fc-NF mini-blocks, to replicate the relationships between the neural and emotional/metacognitive measures as observed in **Experiment 2**, and to assess how neurotransmitter concentrations in NF relevant regions influence these relationships. As in **Experiment 1** and **2**, the number of fc-NF volumes within the correlation window was negatively related with the average fc at each volume (**Fig. 2c**) and positively related with the frequency of negative partial correlations (**Fig. 2d**). We further found that these two measures, i.e. average fc and the frequency of negative partial correlations, plateaued when the fc-NF was estimated consecutively from volumes stemming from fc-NF mini-blocks only (i.e. the ratio of fc-NF : no-NF volumes in the correlation window was 20:0).

Similar to **Experiment 2**, no practice-related change in emotional/metacognitive measures (**Supplementary Table 5**) or practice-related change in fc could be observed at group level (*M* = -.006, *SD* = .061, *p* > .01). Ten individuals showed a negative slope (i.e. change in the desired direction, *M* = -.057, *SD* = .022), nine individuals exhibited a positive slope (*M* = .052, *SD* = .026).

#### Correlation analysis

We first computed the correlation coefficients for the relationships between the neural and emotional/metacognitive measures that were found to be significant in **Experiment 2**. Again, all four correlations were found to be significant. Practice-related change in fc was negatively related with change in thought control ability (TCAQ; **Fig. 3a** bottom; *r*_(17)_ = -.57, B_10_ = 6.04; after the removal of two outliers: *r*_(15)_ = -.66, B_10_ = 13.27). Initial fc was positively related with trait anxiety (STAIT T; **Fig. 3b** bottom; *r*_(17)_ = .43, B_10_ = 1.32; after the removal of one outlier: *r*_(16)_ = .56, B_10_ = 4.31) and negatively related with thought control ability (TCAQ: **Fig. 3c** bottom; *r*_(17)_ = -.55, B_10_ = 4.61; after the removal of three outliers: *r* _(14)_ = -.69, B_10_ = 16.54). The effect of fc-NF was negatively related to state anxiety (STAI-S; **Fig. 3d** bottom; *r*_(17)_ = -.69, B_10_ = 40.49; after the removal of three outliers: *r*_(14)_ = -.80, B_10_ = 170.20).

In addition, **Fig. 4** shows the correlation matrix for the relevant emotional/metacognitive measures, neural measures and neurotransmitter concentrations.

**Fig. 4.**
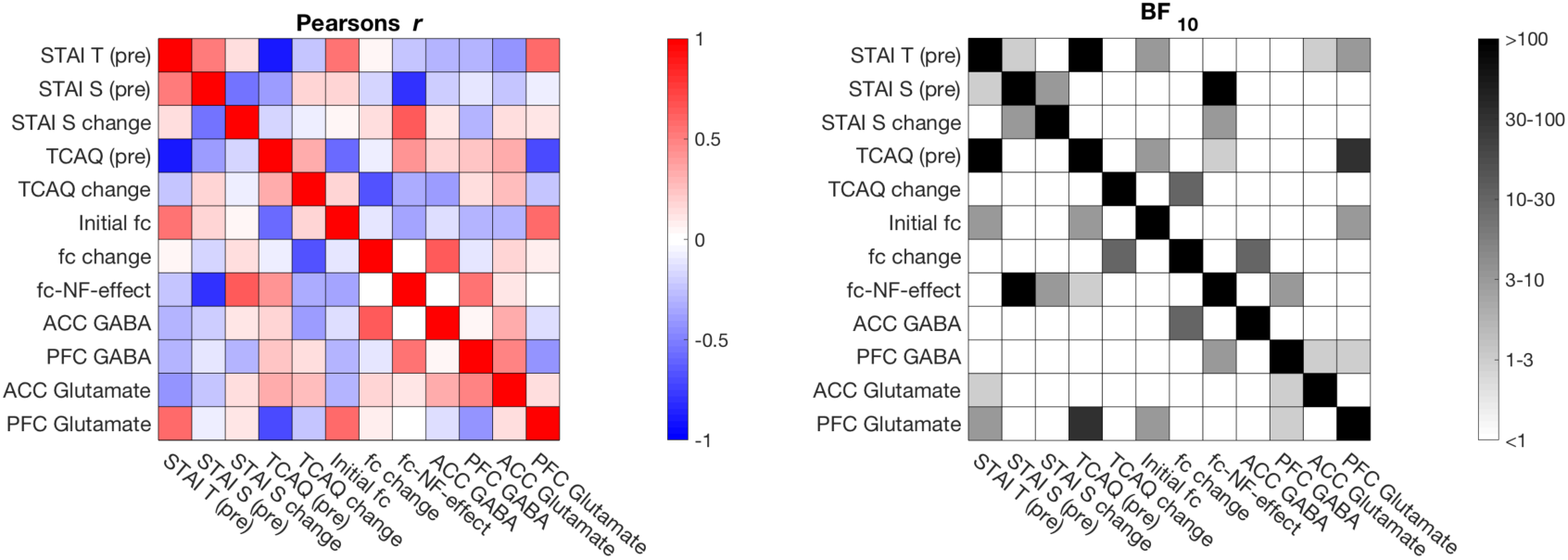
Correlation matrix for relevant emotional/metacognitive measures, neural measures and neurotransmitter concentrations. Bayesian correlation values are shown after outlier removal using Mahalanobis distance.

#### Moderation analysis

We conducted moderation analyses to assess if neurotransmitter concentrations, i.e. glutamate and GABA, in relevant regions influence the significant relationships found between neural measures and emotional/metacognitive measures. PFC GABA and ACC GABA concentrations moderate the relationship between state anxiety before the MRI session and fc-NF-effect (model 3, *F*_(7,11)_ = 3.65, *p* = .028, *R*^*2*^ = .70, **Fig. 5a**). We found STAI-S (*b* = -.009 *t*_(11)_ = −3.514, *p* = .005, 95% CI [-.015, -.003]) and PFC GABA concentrations (*b* = .036 *t*_(11)_ = 2.635, *p* = .023, 95% CI [.006, .066]) to be significant predictors as well as the interaction term between PFC GABA and ACC GABA concentrations to be significant (*b* = -.037 *t*_(11)_ = −2.394, *p* = .036, 95% CI [-.071, -.003], **Table 1**).

To further explore the nature of the interaction we performed a simple slopes analysis. We found that the relationship between STAI-S and the fc-NF-effect reached significance at medium ACC GABA and medium PFC GABA concentrations (*b* = -.009 *t*_(11)_ = −3.513, *p* = .005, 95% CI [-.015, -.003]) and is at trend level for medium ACC GABA and low PFC GABA concentrations (*b* = -.022, *t*_(11)_ = −1.894, *p* = .085, 95% CI [-.048, .004]), whereas no such effect was found at medium ACC GABA and high PFC GABA concentrations (*b* = .004 *t*_(11)_ = .324, *p* = .752, 95% CI [-.023, .031]). No significant effects were observed for low and high ACC GABA (all *p*’s > .1). To test the specificity of this effect we ran the same model using glutamate only (i.e. PFC glutamate and ACC glutamate concentrations) and a combination of GABA and glutamate (e.g. PFC GABA and ACC glutamate concentrations). In none of these cases the overall model was found to be significant (all *p*’s > .1).

**Table 1.** Estimating the influence of neurotransmitter concentrations in NF relevant regions on the relationship between state anxiety before the MRI session and fc-NF-effect.

|  |  | Coeff | SE | t | p | LLCI | ULCI |
| --- | --- | --- | --- | --- | --- | --- | --- |
| Constant | $i_Y$ | -.028 | .020 | -1.438 | .178 | -.072 | .015 |
| STAI-S (X) | $b_1$ | -.009 | .003 | -3.514 | **.005** | -.015 | -.003 |
| PFC GABA (W) | $b_2$ | .036 | .014 | 2.635 | **.023** | .006 | .066 |
| XW | $b_3$ | .006 | .005 | 1.121 | .286 | -.006 | .017 |
| ACC GABA (Z) | $b_4$ | -.029 | .025 | -1.185 | .261 | -.084 | .025 |
| XZ | $b_5$ | -.001 | .005 | -.233 | .820 | -.011 | .009 |
| WZ | $b_6$ | -.037 | .015 | -2.394 | **.036** | -.071 | -.003 |
| XWZ | $b_7$ | .002 | .008 | -.231 | .822 | -.018 | .015 |

**Fig. 5.**
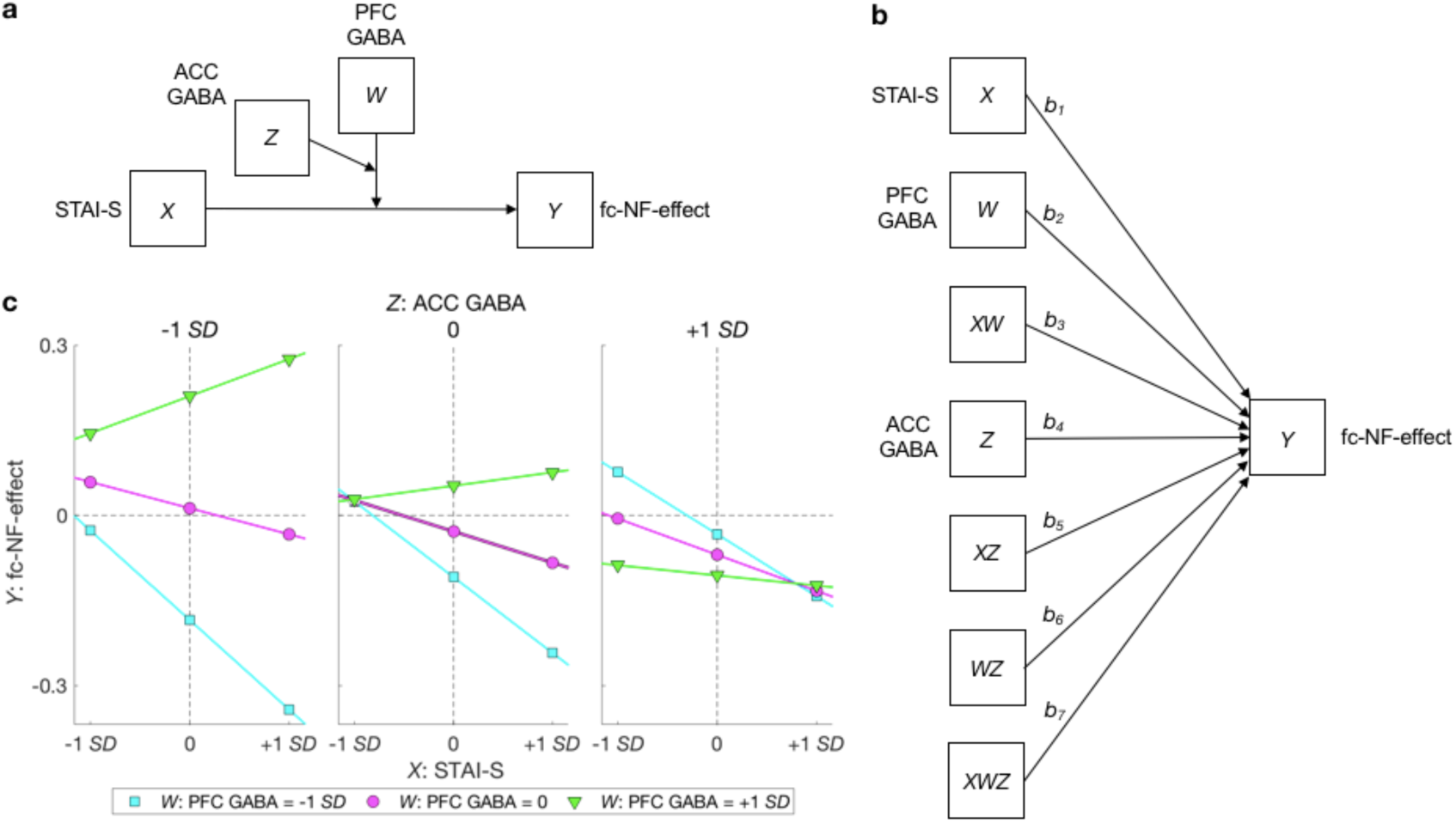
Moderation model and results. Moderation of the effect of state anxiety before the MRI session on the fc-NF-effect by neurotransmitter concentrations in NF relevant regions, depicted as a conceptual diagram (**a**) and a statistical diagram (**b**). (**c**) Simple slopes for STAI-S predicting fc-NF-effect for different levels (+1 *SD*, 0, −1 *SD*) of PFC GABA and ACC GABA concentrations.

## 4. DISCUSSION

Here we aimed at comparing different NF implementations with regard to their effectiveness in modulating PFC-amygdala fc (**Experiment 1**) and at assessing the effects of the most effective NF implementation on neural measures and emotional/metacognitive measures, and their associations (**Experiment 2** and **3**). We further assessed the effect of a longer fc-NF block and whether neurotransmitter concentrations in NF relevant regions influence fc-NF-effects or practice-related changes (**Experiment 3**).

### Differential effects of NF implementations

In **Experiment 1**, we compared three NF implementations and found that the weighted negative NF implementation was most effective in achieving a more negative fc when comparing fc-NF to no-NF. Considering that younger individuals and anxious individuals of all ages exhibit generally lower levels of negative PFC-amygdala fc, or even positive PFC-amygdala fc, it is plausible to suggest that a NF implementation ranging from zero (no correlation) to minus one (maximum negative correlation) was too unspecific to benefit the individuals (Hattie and Timperley, 2007; Lotte et al., 2013). In other words, the naturally more rarely occurring negative connectivity pattern does not receive enough reinforcement to allow the individuals to learn and improve. This finding holds important implications for the development of NF interventions, as it suggests that individually tailored NF implementations should enhance the modulation of task-specific activity patterns even more.

### Absence of significant effects at group level

In **Experiment 2** and **3** we did not find significant changes in emotional/metacognitive measures or in the change in fc at group level. This could be due to several reasons, such as the large inter-individual differences in neural and emotional/metacognitive measures, or the relatively short practice time. We note though that the relationship between the amount of practice and manifestation of emotional/metacognitive changes is not yet clear, and may depend on various aspects (e.g. population, task, NF implementation). While Zilverstand and colleagues (Zilverstand et al., 2015) showed clinically relevant effects with only one NF session, others argue that multiple sessions may be necessary (Linden et al., 2012; Scheinost et al., 2013). For one, this shows that there is an urgent need to actively research the optimal dose of NF practice. Moreover, it may be beneficial to link the NF practice more closely to cognitive-behavioural approaches, such as cognitive behavioural therapy.

### Relationships between emotional/metacognitive and neural measures

The significant associations between emotional/metacognitive and neural measures that we observed in **Experiment 2** were replicated in an independent sample in **Experiment 3**. The observed correlational results can be subdivided into effects of change and predictive effects. Correlations representing effects of change demonstrated that practice-related change in fc is related to change in thought control abilities. Effects of change show that we can use fc-NF not only to selectively modulate specific brain network connections, but also that these neural changes relate to changes in emotional/metacognitive measures, which further supports the effectiveness of the NF implementation.

Results depicting predictive effects found that lower trait anxiety predicted more negative fc. This finding is consistent with previous studies (Delli Pizzi et al., 2017; Kim et al., 2011). Our results further indicate that thought control abilities were negatively related to the initial fc, i.e. the better the thought control ability, the more negative the initial fc. This is not surprising given the negative relationship between trait anxiety and thought control abilities. Together, these results highlight that levels of anxiety and thought control abilities can be used to predict PFC-amygdala fc during emotion regulation.

Our results also provide some insights into individual differences in fc-NF-effects. Specifically, we found that state anxiety prior to the MRI session was negatively related with the fc-NF-effect, i.e. the lower the state anxiety, the higher the effect of fc-NF on fc. This highlights the importance of an individual’s state prior to the NF. Future studies aiming at increasing NF success should therefore ensure that individuals, especially those with high levels of state anxiety, are relaxed and comfortable prior to the NF session. It is fair to assume that an individual’s state not only plays a pivotal role prior to the NF session, but also during the NF session. One approach could therefore be to only initiate the next trial if the individual is in a ‘good’ state (i.e. relaxed, low state anxiety), which could be assessed either behaviourally or by means of an identified a biomarker (Meinel et al., 2016).

### Moderation through neurotransmitter concentrations

In **Experiment 3**, we extended our protocol to obtain glutamate and GABA concentrations from the PFC and ACC. We choose the PFC due to the key role of γ-aminobutyric acid (GABA)-ergic neurotransmission within the PFC in balancing the amygdala activity (Constantinidis et al., 2002), whereas the ACC was chosen due to its centrality for explicit processing of reward, such as NF (Emmert et al., 2016; Sitaram et al., 2016). Our results showed that the relationship between state anxiety (STAI-S) before the MRI session and fc-NF-effect (**Fig. 3d**) is moderated by GABA concentrations in the ACC and PFC. Simple slope analysis revealed that the relationship between STAI-S and the fc-NF-effect reached significance at medium ACC GABA and medium PFC GABA concentrations and is at trend level for medium ACC GABA and low PFC GABA concentrations. This demonstrates that medium ACC GABA concentrations constitutes an optimal level of inhibition. Due to the central role of the ACC for explicit reward processing (Emmert et al., 2016; Sitaram et al., 2016), this result suggests that medium inhibition levels in the ACC are ideal for explicit reward processing, such as NF. Our results further demonstrate that medium and low PFC GABA concentrations, but not high PFC GABA concentrations, promote the relationship between STAI-S and fc-NF-effect. This is in agreement with animal studies demonstrating that higher GABA concentrations in the ventromedial PFC reduces GABA-ergic inhibition of the amygdala, promoting its hyperactivity (Akirav and Maroun, 2007; Chefer et al., 2011; Courtin et al., 2014; Gauthier and Nuss, 2015). Delli Pizzi and colleagues provided supporting evidence for this in humans, i.e. they found a positive relationship between medial PFC GABA concentrations and resting state ventromedial PFC-amygdala fc (Delli Pizzi et al., 2017). This suggests that if high PFC GABA concentrations imply higher amygdala activity, this could in turn result in a more positive PFC-amygdala fc and consequently reduce fc-NF-effects. In sum, our results provide first empirical evidence for the relevance of neurotransmitter concentrations for the effectiveness of fc-NF.

Given the importance of the optimal neurotransmitter concentrations for fc-NF to be effective approaches to alter neurotransmitter concentrations prior to NF should be explored. However, the use of GABA-ergic drugs during childhood and adolescence is controversial due to the risk of developing dependence and severe adverse effects (Sidorchuk et al., 2018). Yet other means that alter GABA-ergic activity, such as yoga (Streeter et al., 2010, 2007), may be adopted. Such studies would also shed further light on the causal interpretation.

A pressing question is whether the observed moderation effect is specific to the domain of emotion regulation, i.e. domain specific, or universal. Within our moderation model ACC GABA concentrations (*Z*) and fc-NF-effects (*Y*) constitute NF specific variables, whereas STAI-T (*X*) and PFC GABA concentrations (*Z*) are domain specific variables. It is thus reasonable to assume that the model is still valid if *X* and *Z* are replaced by corresponding variables in the different domain, such as residual motor function in the affected limb as *X* and GABA concertation from the sensorimotor area of the affected hemisphere as *Z* for motor control following post-stroke. It can further be assumed that this effect may not specific to fc-NF only, but also apply for activity-based NF. Empirical research and computational modelling should be consulted to answer the question wheatear our results are domain specific or universal.

### Implications for translational, clinical application

The results from the current series of experiments have provided strong evidence that fc-NF is feasible in adolescent females with different anxiety levels. We further found that practice-related changes in fc is related to changes in thought control ability. Thus, on a broader scale, our results provide first evidence for a possible clinical application that aims to shape emerging fc patterns non-invasively in the developing brain(Cohen Kadosh et al., 2013). This is significant given that one in four children have increased levels of worry and fear as they enter adolescence and paediatric anxiety predicts lifelong mental disorders (Keshavan et al., 2014). Aberrant emotional regulation strategies arising in anxiety mirror fc patterns in younger individuals where lower levels of negative PFC-amygdala fc or even positive PFC-amygdala fc are exhibited (Kim et al., 2011).

Further research is now necessary to evaluate the effects of multi-session protocols in clinical samples using both subjective and objective clinical outcome measures.

### Limitations

The following limitations merit comment: Despite converging results across the experiments, multi-centre studies are now necessary to examine the effect of psychosocial and socioeconomic factors. Furthermore, we acknowledge that we performed a relatively large number of correlations, whereby we have focussed on priori hypotheses, i.e. correlations with measures of anxiety and thought control ability, and, crucially, used Bayesian inference.

While we underline the acquisition of neurotransmitter levels as a strength of this study, we emphasize that single-voxel MRS has limited spatial resolution. Recent advances in magnetic resonance spectroscopic imaging (MRSI) have the potential to overcome the shortcomings related to single-voxel MRS in future (Steel et al., 2018).

### Conclusions

Our results showed that NF implementations differentially modulate PFC-amygdala fc. Using the most effective NF implementation in a larger sample yielded important associations between neural measures and emotional/metacognitive measures, such as practice-related change in fc was related with change in thought control ability. Further, we found that the relationship between state anxiety and the effect of fc-NF was moderated by GABA concentrations in the PFC and ACC. Future studies that investigate the effects of multi-session protocols in clinical samples and whether the moderation model is domain specific or universal are recommended.

## ACKNOWLEDGEMENTS

We thank Dr. Ralf Veit for discussions regarding the offline MRI data analysis, Prof. Roi Cohen Kadosh for discussions regarding the moderation analysis and Prof. David Linden for comments on the manuscript.

We thank Verena Koppe for her valuable contribution to data management as well as Samantha Clarkstone, Kerenza Hood, Rachel McNamara, Rebecca Playle, Vincent Poile, Elizabeth Randell and Gareth Watson (Centre for Trials Research, College of Biomedical & Life Sciences, Cardiff University, UK), and all BRAINTRAIN coordinators and collaborators for helpful and inspiring discussions.

This work has received funding from the European Union’s Seventh Framework Programme for research, technological development and demonstration under grant agreement no. 602186 and registered as preclinical trial #NCT02463136.

## AUTHOR CONTRIBUTIONS

CZ, ML, SPWH, JL and KCK designed experiment 1 and 2. CZ and KCK designed experiment 3. CZ, NJ, SL and KCK collected the data. NJ analysed the MRS data, CZ performed all other data analyses. CZ and KCK wrote the manuscript and all of the other authors edited the manuscript.

## Supplementary Materials

### Supplementary Methods

#### MRI data acquisition

Before the experimental tasks a high-resolution structural scan was obtained from each subject using a T1-weighted magnetization-prepared rapid-acquisition gradient echo (MPRAGE) sequence (TR = 1900 ms, TE = 3.97 ms, filp angle = 8°, slice thickness 1 mm, in-plane resolution 1 x 1 mm, orientation = sagittal). Functional data were recorded using 2D multiband gradient echo planer imaging (Todd et al., 2016) (TR = 933 ms, TE = 33.40 ms, flip angle = 64°, slice thickness 2 mm, in-plane resolution 2 x 2 mm, orientation = transversal, 72 slices, multi-band factor = 6).

For **Experiment 1** and **2** the localizer comprised 570 volumes and each of the four NF runs comprised 310 volumes. For **Experiment 3** the localizer comprised 300 volumes and each of the three NF runs comprised 330 volumes. At the beginning of the localizer and each NF run ten additional volumes were acquired, but not analysed, to avoid saturation effects.

For **Experiment 3** we collected MRS data. MRS data were acquired using a semi-LASER sequence (Scheenen et al., 2008) (32 averages, TR = 3500 ms, TE = 28 ms, voxel size = 20 × 20 × 20 mm) using VAPOR (variable power RF pulses with optimized relaxation delays) water suppression (Tkác et al., 1999). Two voxels of interest (VOI) were manually positioned. One VOI was positioned at the same position as the PFC ROI, i.e. local maximum of the GLM t-statistics of the brain activity during the localizer (the sum of the three contrasts: appraisal > fixation, reappraisal > fixation, reappraisal > appraisal) within the left dorsolateral and medial PFC. The other VOI, the anterior cingulate cortex (ACC) VOI, was centred on the interhemispheric fissure on the coronal plane superior to the frontal aspect of the corpus callosum on the sagittal plane.

#### Neurofeedback task – Instructions

All participants were blind to their NF implementation and were instructed as follows: *“You will see a thermometer with a green rim on the screen. The red bars show how much the regions that are important for emotions are active. Your job is to get these regions as active as possible! So try to get this thermometer up as much as possible. Similar to the task before, try to control your thoughts towards a positive feeling. When the thermometer does not have a green rim, the thermometer is not working. However, even if the thermometer is not working, your task will be the same and we are still measuring how much your brain is active. The two different thermometers will alternate.*

*When the thermometer has a green rim, it might need some time to properly estimate the brain activity. This leads to a delay in the thermometer display.”* Debriefing revealed that, in order to change their thoughts and emotions towards a happier mood, participants thought about certain people, things that happened in the past and things that they would like to happen in the future.

### Supplementary Results

#### NF implementations differentially modulate the frequency of negative PFC-amygdala partial correlations

The observed differences in fc between fc-NF and no-NF as illustrated in Supplementary Fig. 1 are most likely due to changes in the frequency of negative partial correlations (secondary measure, **Supplementary Fig. 2a**).

The relationship of fc-NF and fc was assessed by correlating the frequency of negative partial correlations at each of the 20 volumes with the number of fc-NF volumes within the correlation window (**Supplementary Fig. 2b**). For the negative NF implementation and the positive NF implementation, the frequency of negative partial correlations and the number of fc-NF volumes in the correlation window were related negatively at trend level (negative NF implementation: *r*_(38)_ = -.31, *p* = .05; positive NF implementation: *r*_(38)_ = -.32, *p* = .05). Contrary, for the weighted negative NF implementation a positive relationship was observed (*r*_(38)_ = .47, *p* < .01), indicating that a higher number of fc-NF volumes within the correlation window equalled a higher frequency of negative partial correlation.

The relationship between the frequency of negative partial correlations and the number of fc-NF volumes in the correlation window was significantly different for the weighted negative NF implementation in comparison to the other two NF implementations (both *p*’s < .001), whereby the other two NF implementations did not differ regarding this aspect (*p* = .96).

#### NF implementations differentially modulate the frequency of correlation scenarios

One limitation of using a fc-NF approach that is based on correlational measures is that we cannot directly infer the direction of change within each region. In other words, a negative correlation exits when variable 1 increases and variable 2 decreases, or when variable 1 decreases and variable 2 increases. However, given the top-down PFC regulation of amygdala reactivity the main goal of PFC-amygdala fc-NF constitutes PFC increase and amygdala decrease.

To understand the nature of the results depicted in Supplementary Fig. 1 and 2, we assigned each negative correlation to one out of four scenarios (see **Supplementary Fig. 5** for details). **Supplementary Fig. 3a** illustrates the percentage of each scenario for fc-NF and no-NF mini-blocks.

As above, we further investigated the relationship between the number of fc-NF volumes within the correlation window and the frequency of scenario 1, i.e. PFC up-regulation & amygdala down-regulation, and of scenario 2, i.e. PFC down-regulation & amygdala up-regulation, at each volume (**Supplementary Fig. 3b**). For the negative NF implementation and the weighted negative NF implementation the number of fc-NF volumes in the correlation window was positively related to the frequency of scenario 1 (negative NF implementation: *r*_(38)_ = 0.39, *p* = .01; weighted negative NF implementation: *r*_(38)_ = .70, *p* < .001) and negatively related to the frequency of scenario 2 (negative NF implementation: *r*_(38)_ = -.50, *p* < .01; weighted negative NF implementation: *r*_(38)_ = -.64, *p* < .001). In contrast, for the positive NF implementation the pattern was reversed, i.e. the number of fc-NF volumes in the correlation window was negatively related to the frequency of scenario 1 (*r*_(38)_ = -.44, *p* < .01) and positively related to the frequency of scenario 2 (*r*_(38)_ = .55, *p* < .001). This suggests that only NF implementations reinforcing negative PFC-amygdala fc support the task-relevant negative fc pattern.

Further, we compared the two negative NF implementations with regard to the strength of the relationship between the number of fc-NF volumes in the correlation window and the frequency of scenario 1. We found a significant difference, indicating that the relationship is strongest for the weighted negative NF implementation.

### Supplementary Discussion

#### Length of the correlation window

We chose a window length of 20 volumes based on a previous study by Zilverstand and colleagues (Zilverstand et al., 2014), who found that the fc measures in a well-controlled, simple motor task were weaker and less reliable for a shorter (12 s) window but not qualitatively different form a longer (26 s) window. Given the higher reliability of the longer correlation windows, we matched the size of our time-windows with the length of the mini-blocks. This allowed us to directly assess the effect of fc-NF as a function of the ratio of fc-NF : no-NF volumes in the correlation window. We found that the PFC-amygdala fc (Supplementary Fig. 1, Fig. 2a,c), frequency of negative partial correlations (Supplementary Fig. 2, Fig. 2b,d), and frequency of scenarios describing different patterns of activation changes (Supplementary Fig. 3) changed as a function of fc-NF volumes within a correlation window. This finding is not only of theoretical interest, but also has important practical implications. Specifically, it suggests that with regards to clinical applications the relation between task window and correlation window should be adapted, so that the length of the task window exceeds the correlation window, in order to further enhance the fc-NF-effect and practice-related changes.

### Supplementary Figures

**Supplementary Fig. 1.**
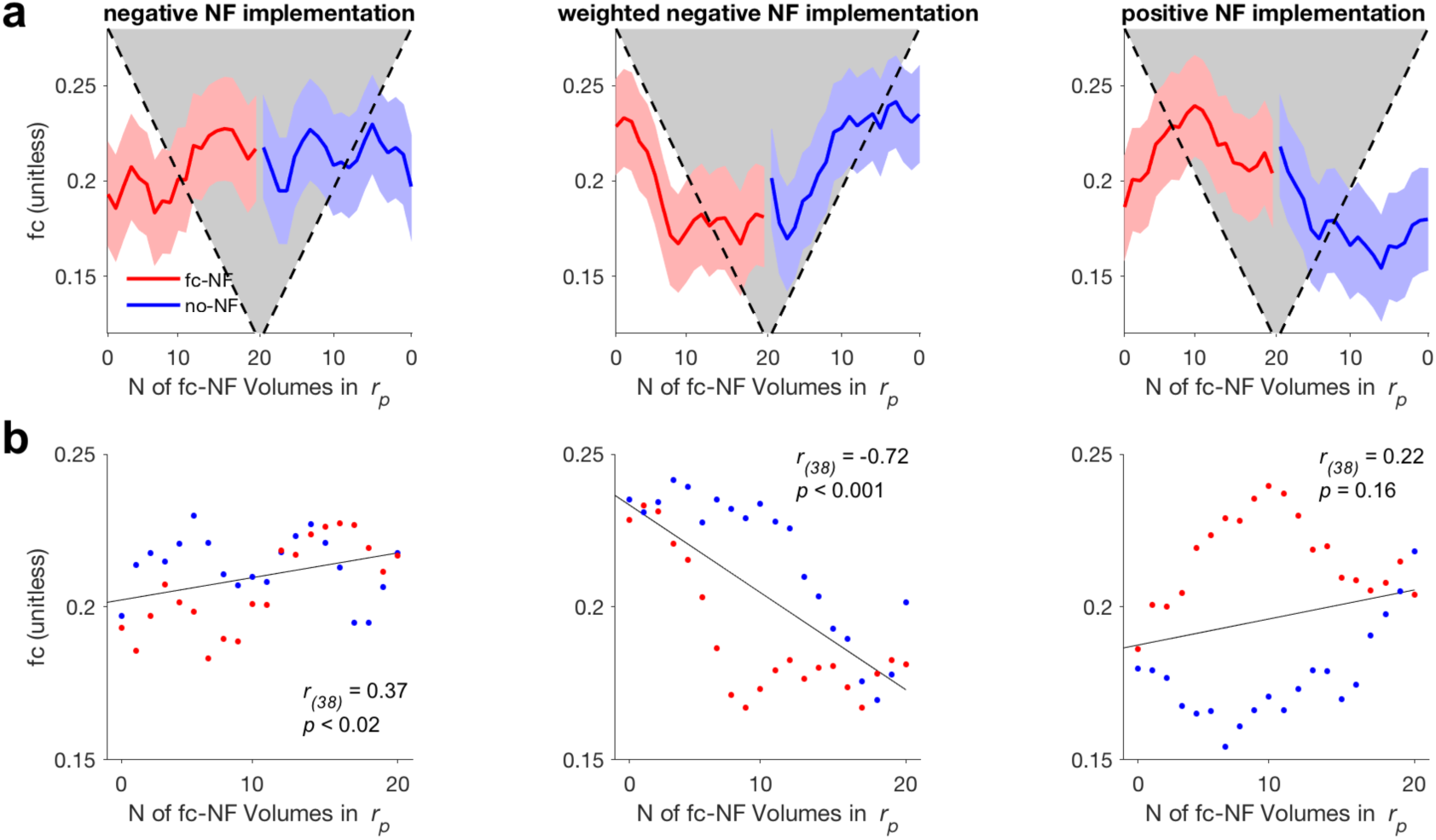
Difference in average functional connectivity (fc) between fc-NF (red) and the no-NF (blue) for each NF implementation (*N* = 5 per NF implementation). **(a)** Means and standard error for the average fc, i.e. partial correlations *r*_*p*_, at each volume within a mini-block. Fc is averaged across mini-blocks within one condition (fc-NF, no-NF), NF runs (1, 2, 3, 4) and individuals within one NF implementation (negative NF implementation, weighted negative NF implementation, positive NF implementation). Number of fc-NF volumes in the correlation window (N of fc-NF Volumes in *r*_*p*_) is illustrated in form of a grey triangle. **(b)** Relationship between average fc at each volume and the number of fc-NF volumes within the correlation window.

**Supplementary Fig. 2.**
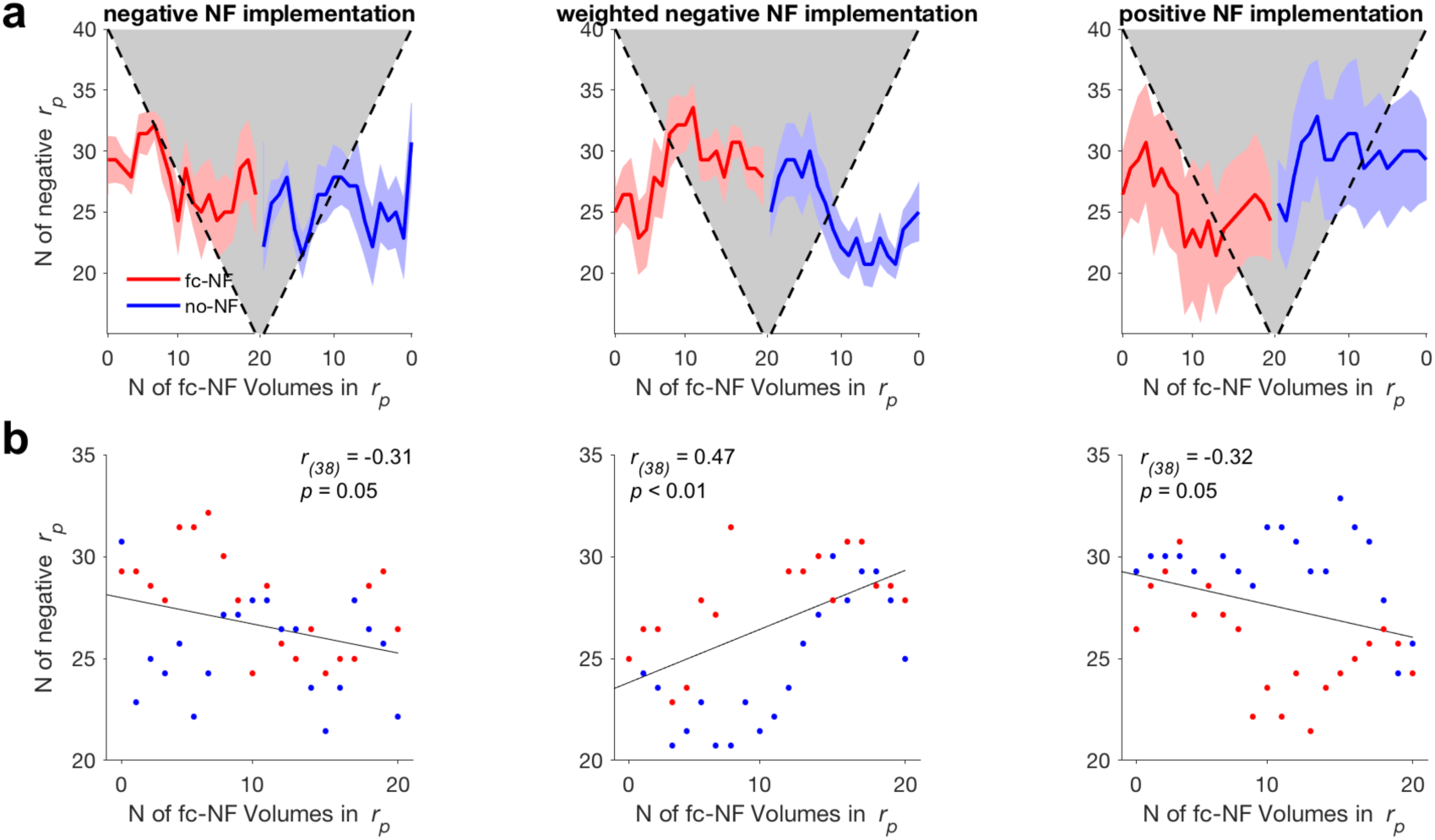
As Supplementary Fig. 1, but for the frequency of negative partial correlations (N of negative *r*_*p*_).

**Supplementary Fig. 3.**
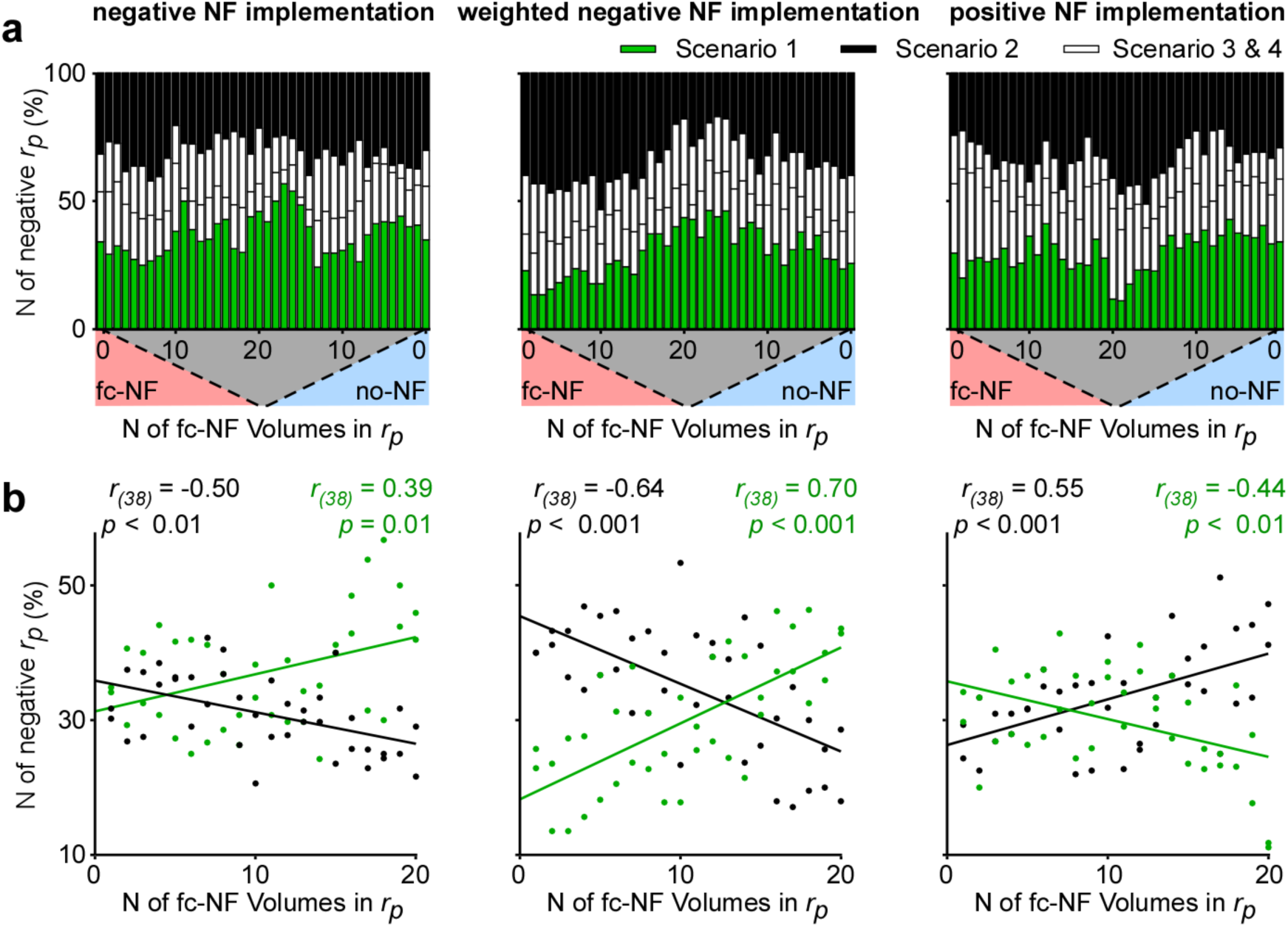
Direction of activity change within the negative partial correlations for each NF implementation (*N* = 5 per implementation). **(a)** Percentage of negative partial correlations between PFC and amygdala for scenario 1 (PFC up-regulation & amygdala down-regulation; green), scenario 2 (PFC down-regulation & amygdala up-regulation; black) or scenario 3 and 4 (white). Percentages are averaged across mini-blocks within one condition (fc-NF, no-NF), NF runs (1, 2, 3, 4) and individuals within one NF implementation (negative NF implementation, weighted negative NF implementation, positive NF implementation). The number of fc-NF volumes in the correlation window is illustrated in form of a grey triangle. **(b)** Relationships between the number of fc-NF volumes within the correlation window (N of fc-NF Volumes in *r*_*p*_) and frequency of scenario 1 (PFC up-regulation & amygdala down-regulation; green). or scenario 2 (PFC down-regulation & amygdala up-regulation; black)

**Supplementary Fig. 4.**
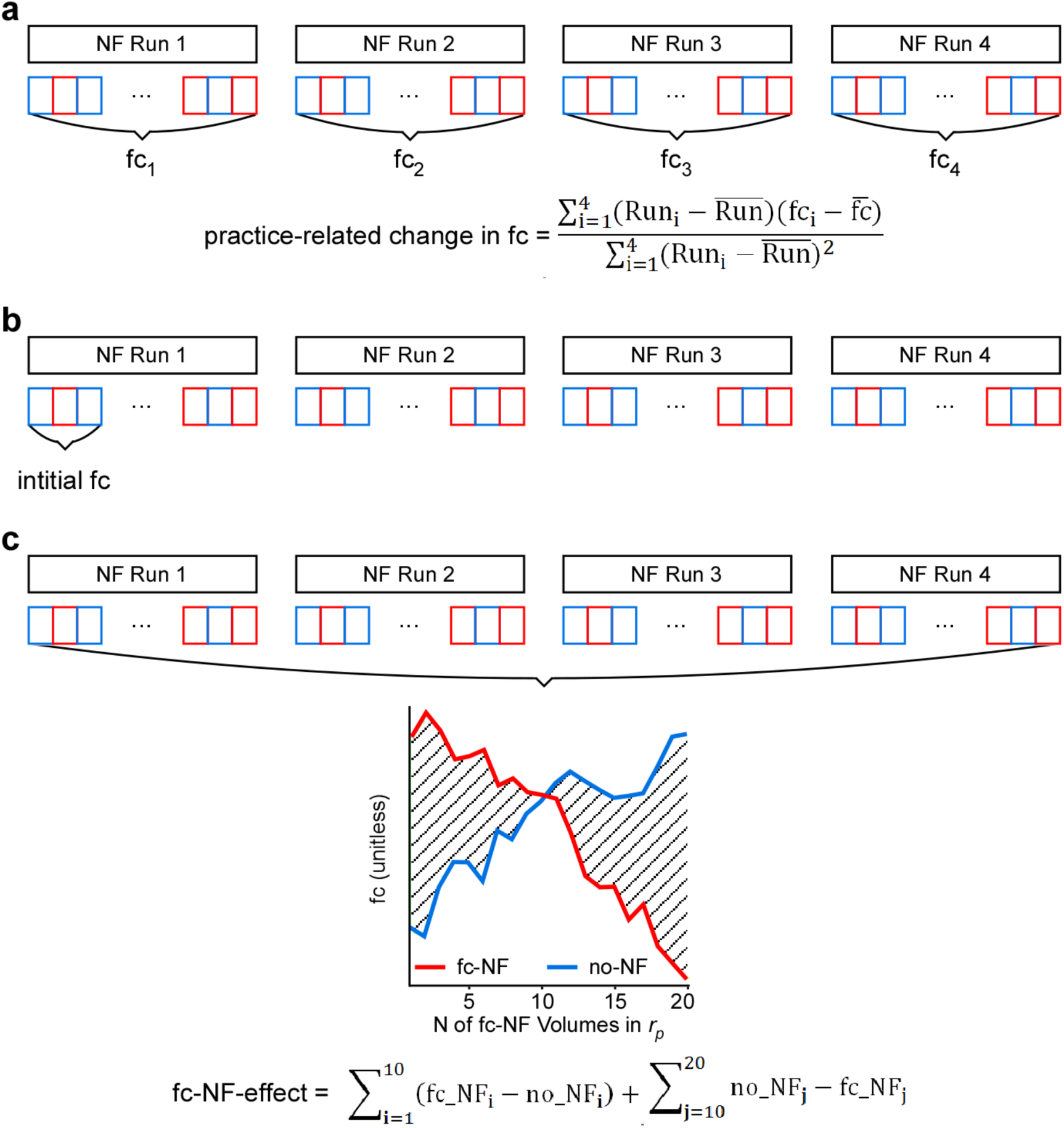
Illustration of quantification of neural measures. (**a**) Visualization of practice-related change in fc, i.e. slope of the linear regression for the average fc within each NF run. (**b**) Illustration of initial fc, i.e. average fc of the first two mini-blocks. (**c**) Visualization of the fc-NF-effect (hatched) on exemplary data from one subject. Due to the dynamic character of the fc-NF, the fc-NF-effect is composed of two parts. For fc-NF mini-bocks (red) the ratio of fc-NF : no-NF volumes in the correlation window is 1:19 at the first volume of the mini-block, 2:18 at the second volume of the mini-block and 20:0 at the last volume of the mini-block. Consequently, in fc-NF mini-blocks volumes 1 to 10 are dominated by no-NF volumes, and volumes 11 to 20 are dominated by fc-NF volumes. Exactly the opposite is the case for no-NF mini-blocks.

**Supplementary Fig. 5.**
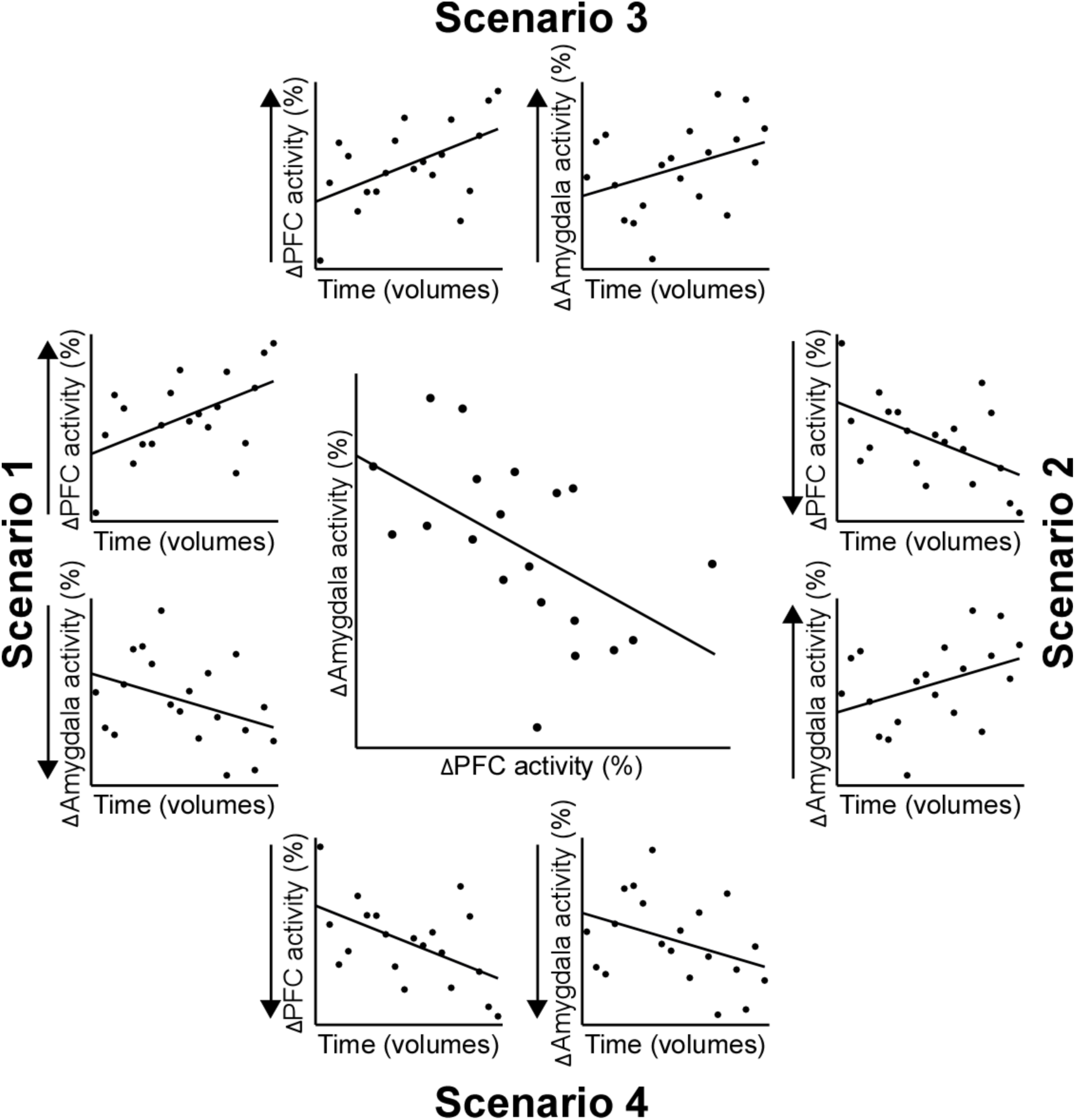
Schematic representation of the procedure used to quantify the direction of activity change within a negative partial correlation. **Centre**: Exemplary negative partial correlation between PFC and amygdala activity accounting for the ‘activity’ in the CST. The direction of activity change within this correlation was determined for the PFC and the amygdala separately. In detail, both, the PFC and the amygdala, were correlated separately with time while accounting for CST ‘activity’. Theoretically four different scenarios are possible: scenario 1 – PFC up-regulation & amygdala down-regulation, scenario 2 – PFC down-regulation & amygdala up-regulation, scenario 3 – PFC & amygdala up-regulation and scenario 4 – PFC & amygdala down-regulation.

### Supplementary Tables

**Supplementary Table 1.** Emotional/metacognitive measures for **Experiment 1**.

| *Pre* | *Negative NF*<br>*implementation*<br>*(N=5)* | *Weighted negative NF*<br>*implementation (N=5)* | *Positive NF*<br>*implementation*<br>*(N=5)* |
| --- | --- | --- | --- |
| *CERQ<sub>pre</sub>* | *M = 59.50; SD = 9.26* | *M = 57.40; SD = 7.33* | *M = 53.65; SD = 10.65* |
| *MFQ<sub>pre</sub>* | *M = 2.60; SD = 2.07* | *M = 1.50; SD = 1.91* | *M = 2.20; SD = 1.71* |
| *SAS-S* | *M = 46.00; SD = 13.41* | *M = 48.00; SD = 11.22* | *M = 56.80; SD = 14.62* |
| *STAI-S<sub>pre</sub>* | *M = 31.40; SD = 8.91* | *M = 23.00; SD = 6.38* | *M = 36.80; SD = 5.22* |
| *STAI-T* | *M = 35.20; SD = 7.92* | *M = 35.80; SD = 8.56* | *M = 38.00; SD = 9.92* |
| *TCQ<sub>pre</sub>* | *M = 65.75; SD = 4.27* | *M = 67.20; SD = 9.26* | *M = 59.60; SD = 7.50* |
| *TCAQ<sub>pre</sub>* | *M = 79.40; SD = 14.62* | *M = 82.00; SD = 10.42* | *M = 75.60; SD = 13.63* |

*CERQ* = Emotion Regulation Questionnaire; *MFQ* = Mood and Feelings Questionnaire; *SAS-S* = Social Anxiety Scale for Adolescents; *STAI-S*, *STAI-T* = State-Trait Anxiety Inventory; *TCQ* = Thought Control Questionnaire; *TCAQ* = Cognitive Thought Control Ability Questionnaire

**Supplementary Table 2.** Emotional/metacognitive measures for **Experiment 2**.

|  | *Pre* | *Post* | *Difference* |
| --- | --- | --- | --- |
| *CERQ* | $N = 27; M = 60.52; SD = 11.09$ | $N = 27; M = 60.93; SD = 11.18$ | $t_{(26)} = -.442; p = .662$ |
| *MFQ* | $N = 25; M = 5.44; SD = 4.81$ | $N = 27; M = 5.56; SD = 4.56$ | $t_{(24)} = -.778; p = .444$ |
| *SAS-S* | $N = 27; M = 49.15; SD = 15.01$ | | |
| *STAI-S* | $N = 24; M = 32.79; SD = 8.61$ | $N = 27; M = 31.78; SD = 9.83$ | $t_{(23)} = 1.784; p = .088$ |
| *STAI-T* | $N = 27; M = 43.52; SD = 11.49$ | | |
| *TCQ* | $N = 27; M = 61.04; SD = 8.67$ | $N = 27; M = 60.70; SD = 8.99$ | $t_{(26)} = .251; p = .804$ |
| *TCAQ* | $N = 23; M = 78.91; SD = 15.37$ | $N = 27; M = 79.85; SD = 14.77$ | $t_{(22)} = .041; p = .968$ |

*CERQ* = Emotion Regulation Questionnaire; *MFQ* = Mood and Feelings Questionnaire; *SAS-S* = Social Anxiety Scale for Adolescents; *STAI-S*, *STAI-T* = State-Trait Anxiety Inventory; *TCQ* = Thought Control Questionnaire; *TCAQ* = Cognitive Thought Control Ability Questionnaire

**Supplementary Table 3:** Emotional/metacognitive measures for a subsample of **Experiment 2** (negative level, i.e. change in the desired direction, *N*=13).

|  | *Pre* | *Post* | *Difference* |
| --- | --- | --- | --- |
| *CERQ* | $N = 13; M = 62.92; SD = 12.43$ | $N = 13; M = 64.15; SD = 12.43$ | $t_{(12)} = -.854; p = .410$ |
| *MFQ* | $N = 12; M = 4.92; SD = 5.00$ | $N = 13; M = 4.92; SD = 4.92$ | $t_{(11)} = -.561; p = .586$ |
| *SAS-S* | $N = 13; M = 52.15; SD = 13.45$ | | |
| *STAI-S* | $N = 13; M = 31.83; SD = 9.04$ | $N = 13; M = 30.85; SD = 7.86$ | $t_{(12)} = .616; p = .551$ |
| *STAI-T* | $N = 13; M = 42.54; SD = 12.14$ | | |
| *TCQ* | $N = 13; M = 62.69; SD = 8.60$ | $N = 13; M = 60.77; SD = 10.69$ | $t_{(12)} = 1.402; p = .133$ |
| *TCAQ* | $N = 11; M = 81.27; SD = 15.38$ | $N = 13; M = 80.54; SD = 15.01$ | $t_{(10)} = 1.258; p = .237$ |

*CERQ* = Emotion Regulation Questionnaire; *MFQ* = Mood and Feelings Questionnaire; *SAS-S* = Social Anxiety Scale for Adolescents; *STAI-S*, *STAI-T* = State-Trait Anxiety Inventory; *TCQ* = Thought Control Questionnaire; *TCAQ* = Cognitive Thought Control Ability Questionnaire

**Supplementary Table 4:** Emotional/metacognitive measures for a subsample of **Experiment 2** (positive level, i.e. change in the undesired direction *N*=14).

|  | *Pre* | *Post* | *Difference* |
| --- | --- | --- | --- |
| *CERQ* | $N = 14; M = 58.29; SD = 9.60$ | $N = 14; M = 57.93; SD = 9.34$ | $t_{(13)} = .302; p = .768$ |
| *MFQ* | $N = 13; M = 5.92; SD = 4.79$ | $N = 14; M = 6.14; SD = 4.29$ | $t_{(12)} = -.519; p = .613$ |
| *SAS-S* | $N = 14; M = 46.36; SD = 16.33$ | | |
| *STAI-S* | $N = 12; M = 33.75; SD = 8.43$ | $N = 14; M = 32.64; SD = 11.59$ | $t_{(11)} = 1.170; p = .115$ |
| *STAI-T* | $N = 14; M = 44.43; SD = 11.24$ | | |
| *TCQ* | $N = 14; M = 59.50; SD = 8.76$ | $N = 14; M = 61.57; SD = 7.38$ | $t_{(13)} = -.965; p = .352$ |
| *TCAQ* | $N = 12; M = 76.75; SD = 15.70$ | $N = 14; M = 79.21; SD = 15.08$ | $t_{(11)} = -1.105; p = .293$ |

*CERQ* = Emotion Regulation Questionnaire; *MFQ* = Mood and Feelings Questionnaire; *SAS-S* = Social Anxiety Scale for Adolescents; *STAI-S*, *STAI-T* = State-Trait Anxiety Inventory; *TCQ* = Thought Control Questionnaire; *TCAQ* = Cognitive Thought Control Ability Questionnaire

**Supplementary Table 5.** Emotional/metacognitive measures for **Experiment 3**.

|  | *Pre* | *Post* | *Difference* |
| --- | --- | --- | --- |
| *CERQ* | $N = 19; M = 58.16; SD = 9.00$ | $N = 19; M = 60.84; SD = 11.08$ | $t_{(18)} = -1.326; p = .202$ |
| *MFQ* | $N = 19; M = 6.21; SD = 5.72$ | $N = 19; M = 6.42; SD = 5.82$ | $t_{(18)} = -.200; p = .844$ |
| *SAS-S* | $N = 19; M = 48.00; SD = 15.37$ | | |
| *STAI-S* | $N = 19; M = 31.42; SD = 6.01$ | $N = 19; M = 29.00; SD = 5.85$ | $t_{(18)} = 1.490; p = .153$ |
| *STAI-T* | $N = 19; M = 42.42; SD = 10.70$ | | |
| *TCQ* | $N = 19; M = 60.42; SD = 7.52$ | $N = 19; M = 62.74; SD = 7.90$ | $t_{(18)} = -1.180; p = .254$ |
| *TCAQ* | $N = 19; M = 74.47; SD = 18.20$ | $N = 19; M = 75.58; SD = 19.27$ | $t_{(18)} = -.045; p = .657$ |

*CERQ* = Emotion Regulation Questionnaire; *MFQ* = Mood and Feelings Questionnaire; *SAS-S* = Social Anxiety Scale for Adolescents; *STAI-S*, *STAI-T* = State-Trait Anxiety Inventory; *TCQ* = Thought Control Questionnaire; *TCAQ* = Cognitive Thought Control Ability Questionnaire

## Footnotes

† Based on pilot data (*N* = 5). Cumulative distribution function of PFC-amygdala fc, i.e. partial correlations *r_p_* between the PFC, amygdala and corticospinal tract.

‡ We note that the current study focused on left-lateralized ROIs as those were reliably activated in all subjects in the localiser task, whereby a lateralization in favour of the left hemisphere is in line with the literature (Ochsner and Gross, 2005).

§ Note that the cognitive and psychological measures were comparable between the three groups (see **Supplementary Table 1**).

